# De-Darwinizing the proteome: the genome as the original germ line

**DOI:** 10.64898/2026.05.06.723269

**Authors:** William C. Ratcliff, Anthony J. Burnetti

## Abstract

A hallmark of evolutionary transitions in individuality is the suppression of lower-level Darwinian dynamics through germ-soma specialization. The RNA world hypothesis posits that early life arose through ribozymes, RNA molecules that functioned as both catalysts and hereditary units, housed within protocells. This dual role is the chief appeal of the RNA world framework, because it makes ribozymes capable of open-ended Darwinian evolution. It is also its chief liability. Each ribozyme is a fully Darwinian entity: it replicates, accumulates mutations via Müller’s ratchet, and can be displaced by selfish variants that replicate faster but contribute less to collective function. We show that between-protocell selection, even when strong, cannot overcome this within-cell evolutionary erosion, particularly as protocells evolve increased complexity by increasing the number of ribozyme types they contain. The origin of a low-copy-number informational RNA genome, translated by an ancient ribosome into protein enzymes, resolves this conflict by producing a functional workforce that is non-heritable. Protein-based enzymes are intrinsically stripped of evolutionary agency, confining heritable variation to the genome and entrenching the protocell as the primary level of Darwinian individuality. The genome, in this view, is not merely a storage device but the original germ line, and its origin marks the point at which cellular life became capable of open-ended growth in complexity.

## Introduction

The RNA world is a leading framework for the origin of life because it is both simple and powerful (1, 2). A ribozyme has heritable sequence information and catalytic function in the same molecule, and is therefore capable of open-ended Darwinian evolution (3). Housed within lipid vesicles (4), populations of ribozymes could form metabolic networks of increasing complexity, with protocell-level selection favoring collectives that grow and divide.

But this strength is also a liability. Each ribozyme is fully Darwinian: it replicates, gains mutations, and responds to selection. As the number of distinct ribozyme types *k* in the network grows, two problems become severe. The first is coordination: stochastic partitioning at cell division can fail to transmit all *k* types to both daughters, killing one or both. This problem is well recognized (5–7). The second is evolutionary erosion, and we regard it as the more fundamental constraint. Each time a ribozyme reproduces, it can acquire novel mutations, most of which are deleterious. If unchecked by protocell-level selection, this would drive terminal loss of function via Muller’s ratchet (8). Worse yet, cheats may arise, which are both less functional and faster replicating, sweeping through the intracellular population at the expense of collective function. This is not merely a theoretical concern. Spiegelman’s classic *in vitro* evolution experiments showed that Q*β* replicase, given an RNA template and free nucleotides, drives the population toward progressively shorter molecules that replicate faster but lose biological function (9). More recent compartmentalized RNA replicator experiments have repeatedly observed parasitic RNAs spontaneously emerging from functional templates and threatening the persistence of cooperative replication unless held in check by spatial or temporal structure (10–13). This is the tragedy of the commons at the molecular level. The parallel to cancer in multicellular organisms is direct: in both cases, lower-level units that gain a replicative advantage at the expense of collective function can proliferate within the higher-level individual, and between-individual selection may be too weak or too slow to contain the damage (14). The vulnerability of cooperative molecular networks to parasitic replicators was recognized early in the context of Eigen’s hypercycle (15, 16), and the stochastic corrector model showed that group selection among protocells can partially counteract it, but only under restrictive conditions (5, 7).

We argue that both problems are resolved by a single innovation: the origin of the singular genome per proto-cell, read out through a primitive ribosome into protein-based enzymes. Although resolved together, these two problems are logically separable. The coordination problem could in principle be solved by physical linkage alone: concatenating ribozyme genes into a single long RNA molecule would ensure co-transmission without requiring translation at all. What linkage alone cannot solve is the problem of evolutionary agency. As long as the functional enzymes are themselves evolutionary units, within-cell Darwinian dynamics will degrade collective function regardless of whether the genes are linked or free. Translation of information from a single genome per protocell resolves this deeper problem, by producing a functional workforce (proteins and non-replicating mRNAs) that cannot write heritable changes back to the informational molecule (the genome). A single heritable genome per protocell thus acts as a germ line: it solves coordination through linkage and conflict through de-Darwinization (Figure 1). That such a division of labor can arise spontaneously from conflicting multilevel selection has been demonstrated in simulation (17, 18); our focus here is on its evolutionary consequences.

**Fig. 1.**
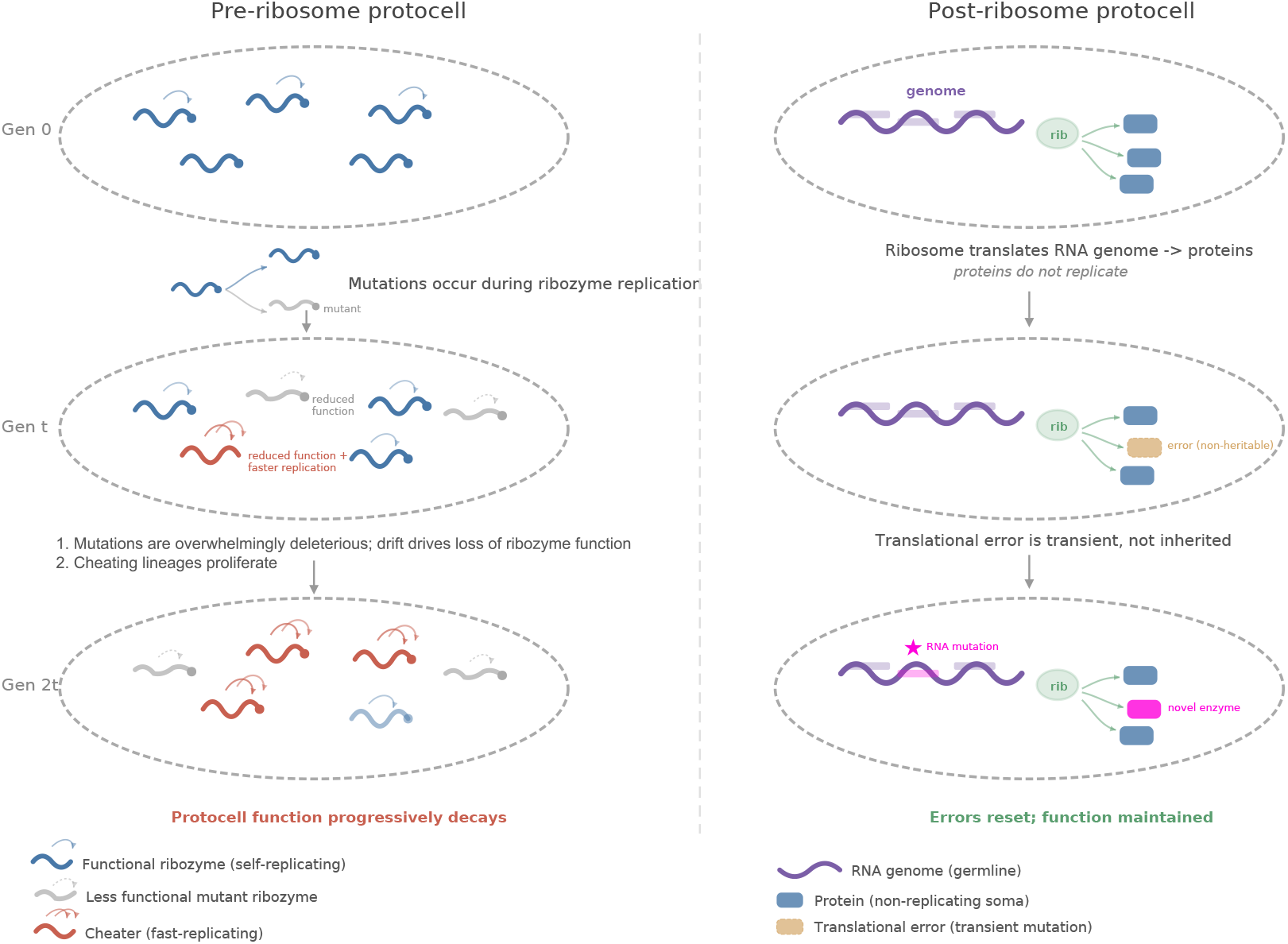
Two architectures of protocell heredity. Left: During ribozyme replication, each has the potential to gain a mutation, which rarely improves function. Mutants are typically neutral, degrade performance (grey), or act as cheats (red) that have both reduced function and higher within-protocell fitness. Protocell function progressively decays. Right: in a post-ribosome protocell, an RNA genome (purple) is translated by a ribosome into protein enzymes (blue). Translational errors (gold) are transient and non-heritable: each generation of proteins is produced fresh from the unchanged genome. Only mutations to the genome itself (pink star) are inherited, and these are subject to protocell-level selection. The genome acts as a germ line; proteins are soma.

## Results

Consider a protocell with *k* ribozyme types, *n* copies each, for a total of *N* = *kn* molecules. At division, each molecule is independently assigned to one daughter with probability 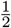, the best-case scenario. For a given type with *n* copies, the probability that one daughter receives none is (1*/*2)^*n*^, and by symmetry the probability that *either* daughter loses that type is 2^1−*n*^. Since types segregate independently:

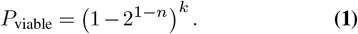

This scales badly (Fig. 2A). With *n* = 5 copies per type: *P*_viable_ ≈ 0.72 for *k* = 5, but only 0.04 for *k* = 50. Maintaining *P*_viable_ ≥ 0.95 at *k* = 50 requires *n* ≈ 11 copies per type, or *N* = 550 total ribozymes, and any variance in copy number across types makes things worse (Fig. 2B). The co-ordination problem imposes a ceiling on network complexity that rises only by increasing total molecular count. Maintaining a fixed viability threshold requires the copy number per type to grow logarithmically with *k*, so the total molecular burden grows faster than linearly (approximately as *k* log *k*). Increased copy number of each molecular type in turn serves to increase the rate of origination of potential cheat mutants, further degrading fitness. A genome collapses these problems to the replication of a single molecule per generation.

**Fig. 2.**
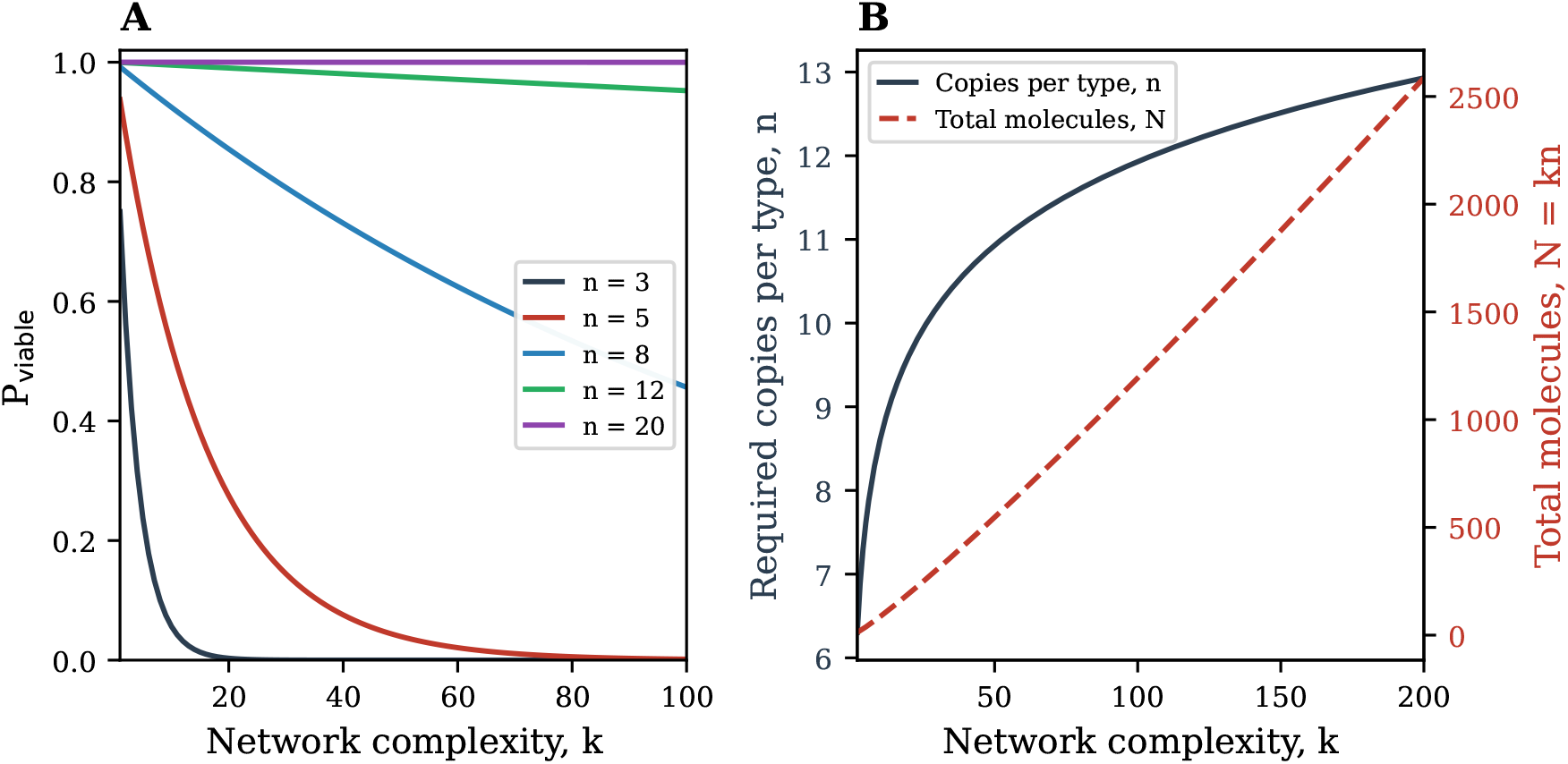
The coordination problem. (**A**) Probability that both daughter protocells inherit at least one copy of all *k* ribozyme types, as a function of network complexity *k*, for different copy numbers per type *n*. Division is symmetric binomial (*p* = 0.5). Even modest network sizes (*k >* 20) require substantial copy numbers to avoid lethal segregation failures. (**B**) Required copy number per type *n* (left axis) and total molecular count *N* = *kn* (right axis) to maintain *P*_viable_ ≥ 0.95. The metabolic burden of maintaining all components grows faster than linearly with network complexity, because the required copy number per type increases logarithmically with *k*.

Physical linkage of genes on a single RNA molecule solves the coordination problem (19), but it does not address evolutionary dynamics within the protocell. Consider a single ribozyme type maintained at *n* copies within a protocell. Each copy replicates, and with probability *µ* per replication, a daughter molecule is degraded or non-functional. We assume that back-mutation is negligible, so functional copies can become degraded but not the reverse. This results in Muller’s ratchet (8): in a finite asexual population, the functional class shrinks irreversibly. Let *x* denote the fraction of functional copies. The expected decline per ribozyme generation is:

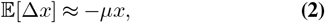

yielding exponential decay 𝔼 [*x*(*t*)] ≈ *e*^−*µt*^. If cheater variants replicate at rate 1+*s* relative to wild-type, within-cell selection accelerates this decline. Across *k* independent types with multiplicative fitness *W* = Π *x*_*i*_, protocell fitness decays as *e*^−*kµt*^ (Fig. 3A), a decline that worsens with every addition to the network. We note that the multiplicative fitness assumption is a simplification: epistatic interactions among ribozyme types could alter the quantitative dynamics. The precise rate of decline depends on the form of the fitness function, but qualitatively similar collapse occurs whenever multiple essential types must each retain function, as the simulations below confirm.

**Fig. 3.**
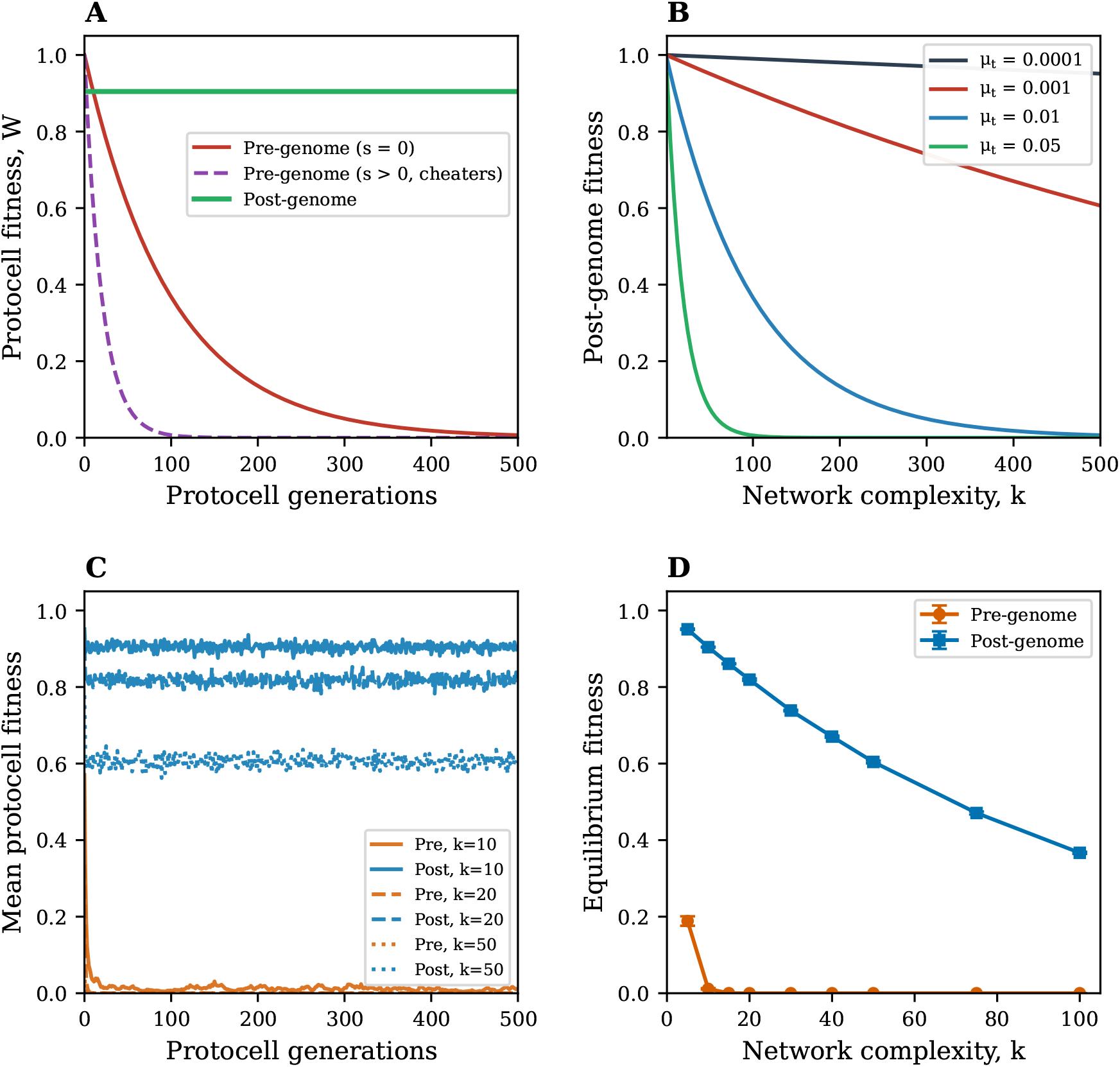
De-Darwinization through translation and multilevel selection. (**A**) Deterministic protocell fitness over generations. Pre-genome fitness decays exponentially as ribozymes accumulate mutations (and faster with cheaters), while post-genome fitness remains constant at (1 − *µ*_*t*_)^*k*^ because translational errors are not inherited. Decay curves use *µ*_eff_ = *µ/*100 for visualization; unscaled dynamics are shown in panels C and D. Parameters: *k* = 20, *µ* = *µ*_*t*_ = 0.005, *τ* = 10, *s* = 0.02. (**B**) Fixed fitness cost of translational error, (1− *µ*_*t*_)^*k*^, as a function of network complexity *k* for different error rates. This cost is constant across generations, not progressive. (**C**) Stochastic simulations with between-protocell selection (*M* = 100 protocells, 500 generations) for pre-genome (orange) and post-genome (blue) architectures at *k* = 10, 20, and 50. Pre-genome protocells decline despite group selection; post-genome protocells maintain stable fitness. Parameters: *µ* = *µ*_*t*_ = 0.005, *s* = 0.02, *τ* = 10, *n* = 10, *U* = 0.005. (**D**) Equilibrium fitness (generations 400–500) as a function of *k*. The pre-genome architecture collapses with increasing complexity; the post-genome architecture remains viable, declining gradually at high *k* as it approaches the genome error threshold. Error bars: s.d. across 5 replicates.

An important simplification throughout this analysis is that all mutations are modeled as deleterious: functional ribozymes can degrade but non-functional variants do not improve. This is a deliberate simplification. The question we ask is whether pre-genome protocells can maintain existing collective function as network complexity grows, not whether they can acquire new capabilities. Even in a system where beneficial mutations occasionally arise, the architecture must first be able to preserve what selection has already built. If within-cell degradation outpaces the capacity of between-cell selection to purge the damage, the system is unsustainable regardless of its adaptive potential.

Between-protocell selection can in principle oppose this, but only if the variance in fitness among protocells exceeds the rate of within-cell degradation. When within-cell dynamics are fast relative to protocell generation time, this condition fails: most protocells converge on similarly degraded states, leaving little variation for selection to act on.

Now consider the post-genome protocell. A singular RNA genome of *k* genes is replicated only once per cell division, with per-gene mutation rate *U* . Each gene is translated into protein, with per-protein translational error rate *µ*_*t*_. We set *µ*_*t*_ = *µ* so that any difference in outcome between the two architectures is attributable to whether errors are heritable, not to their frequency.

In the pre-genome protocell, a mutation in a ribozyme is heritable: the mutant replicates and its lineage carries the change forward. In the post-genome protocell, a translational error in a protein is not: the protein is an informational dead end and eventually degraded, and the next round of translation reads the unchanged genome. Formally, conditional on gene *i* being intact in the genome:

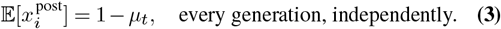

The fraction of functional proteins is reset each generation by translation from the genome. For *µ*_*t*_ = 10^−4^ and *k* = 100, the fitness cost of translational error is (1−*µ*_*t*_)^*k*^ ≈ 0.99: negligible and, crucially, constant (Fig. 3B). A fixed cost of doing business, not a progressive disease.

The post-genome architecture also eliminates within-cell cheating entirely, assuming there are no self-replicating RNA transcripts of individual genes. Proteins cannot replicate as they cannot template their own sequences, so the concept of a faster-replicating variant is meaningless. The only route to increasing the frequency of a non-functional phenotype is to mutate the genome itself, and such a mutation is subject to negative selection at the protocell level. The sign of selection is reversed: from *s >* 0 within the cell (favoring cheaters) to *s*_eff_ *<* 0 between protocells (purging them). Prions, proteins that can template their own conformation, represent a partial exception to this principle (20): by acquiring a form of heritability at the protein level, they partially re-Darwinize the soma. That prions are often pathological underscores how fundamental the suppression of protein-level evolution is to cellular function. Other phenomena such as RNA editing and programmed translational recoding add complexity to the relationship between genome and proteome, but they do not violate the core asymmetry: none of these processes allow a protein variant to rewrite its encoding gene. Transcription of the genome into temporary amplified mRNAs adds an additional error rate to the production of the protein workforce, but unless they are in turn replicated alongside the genome they too do not rewrite the encoding genome. It is again telling that replicating non-genomic RNAs typically represent pathological viruses or selfish transposable elements, working against the fitness of the organism as a whole.

The argument above can be made precise using the Price equation in its multilevel form (21, 22). Consider a population of protocells indexed by *j*, each containing a community of catalytic molecules. Let *Z*_*j*_ denote the mean functional fraction of catalysts in protocell *j*, and let *W*_*j*_ denote the fitness (reproductive output) of protocell *j*, which is an increasing function of *Z*_*j*_. The change in the population-wide mean 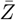 across one protocell generation is:

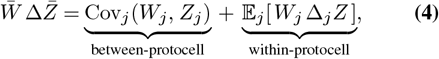

where Δ_*j*_*Z* is the change in mean functional fraction within protocell *j* due to within-cell dynamics during one protocell generation (23).

### Pre-genome protocells

Within each protocell, ribozymes replicate independently. Mutation degrades functional copies at rate *µ* per replication, and cheater variants with replication advantage *s* displace functional types. Over *τ* ribozyme generations per protocell generation, the within-protocell term is negative on average: 𝔼 [Δ_*j*_*Z*] *<* 0, reflecting both mutational degradation and the spread of cheaters. The between-protocell term is positive, since protocell fitness *W*_*j*_ increases with *Z*_*j*_, giving Cov(*W*_*j*_, *Z*_*j*_) *>* 0. The condition for the maintenance of collective function is:

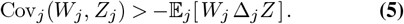

This condition becomes progressively harder to satisfy as network complexity *k* grows, for two reasons. First, within-protocell degradation worsens because mutation and cheater dynamics operate independently across all *k* types, and their effects on protocell fitness are multiplicative (*W* = Π*x*_*i*_), so each additional type amplifies the rate of fitness decline. Second, as within-cell dynamics drive all protocells toward similarly degraded states, the between-protocell variance in *Z*_*j*_ shrinks, reducing the covariance term that selection requires. The simulations described below confirm this dynamic directly. When within-cell generations are fast relative to proto-cell generations (*τ* ≫ 1), the within-protocell term dominates and protocell-level adaptation fails (Fig. 3C).

This is the regime in which the stochastic corrector model (5, 7) operates. That model demonstrated that group selection among protocells can maintain multiple replicator types, but only when the number of types is small, copy numbers are high, and the replicative advantage of cheaters is modest. As Grey et al. (7) showed, even under favorable conditions the stochastic corrector mechanism becomes increasingly fragile as network complexity grows, precisely because the within-protocell selection term in Eq. 5 scales with *k*. Our contribution is to show that translation of non-replicative gene products from a single cellular genome eliminates this term entirely, converting the stochastic corrector’s contingent, parameter-dependent maintenance of cooperation into a robust architectural solution.

### Post-genome protocells

The within-protocell term vanishes when a genome exists. Proteins do not replicate, so there is no within-cell evolutionary process operating on the catalytic workforce. Translational errors are independently and identically distributed each generation, conditioned on the genome, so there is no directional within-cell change: 𝔼 [Δ_*j*_*Z* | genome intact] = 0. The Price equation reduces to:

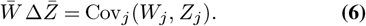

Only between-protocell selection remains. Heritable variation in protocell fitness comes solely from mutations to the genome, which are subject to purifying selection at the protocell level. The genome has eliminated the opposing within-group term by confining heritable variation to a single molecule that does not compete with itself within the cell. This is the formal meaning of de-Darwinization: the within-group component of the Price equation, which drives the tragedy of the commons in the pre-genome architecture, is set to zero by making the amplified catalytic workforce non-heritable.

To test whether between-protocell selection can rescue pre-genome protocells, we simulated a population of *M* = 100 protocells competing over *G* = 500 generations (Fig. 3C,D). In the pre-genome architecture, each protocell contains *k* ribozyme types with *n* = 10 copies each, subject to mutation (*µ* = 0.005 per replication) and cheater advantage (*s* = 0.02) over *τ* = 10 within-cell replication rounds per protocell generation. Crucially, the simulation assumes that the coordination problem has already been solved: daughter protocells inherit the exact ribozyme composition of their parent without stochastic partitioning at division, so that no ribozyme type is lost through segregation. This is a deliberately favorable assumption for the pre-genome architecture, because in reality stochastic loss of entire ribozyme types at division would compound the evolutionary degradation. Even under this best-case scenario, evolutionary erosion alone is sufficient to collapse collective function. In the post-genome architecture, each protocell carries a genome of *k* genes, each subject to irreversible mutation at rate *U* = 0.005 per gene per generation, with translational error rate *µ*_*t*_ = 0.005. Back-mutation is neglected, consistent with the analytical model; the sole mechanism removing deleterious mutations from the population is purifying selection at the proto-cell level. Mutated genes contribute a small residual function *δ* = 0.01 rather than zero, so that protocells carrying mutated genes retain non-zero fitness on which selection can operate. Protocell fitness is the product of per-gene contributions, where each intact gene contributes (1−*µ*_*t*_) in expectation and each mutated gene contributes *δ*. Between-protocell selection operates by differential reproduction proportional to fitness, maintaining constant population size. The per-gene mutation rate *U* is chosen so that the population approaches the genome-level error threshold as *k* grows: the expected number of new deleterious mutations per genome per generation, *kU*, reaches unity at *k* = 200, and the onset of this decline is already visible at *k* = 100 in Fig. 3D, motivating the discussion of DNA below. Even with between-protocell selection, pre-genome fitness declines rapidly as *k* increases, because within-cell degradation erodes the between-protocell variance that selection requires (Fig. 3C). Post-genome protocells, by contrast, maintain high fitness across a wide range of *k*, losing function only when per-gene mutation rates are high enough to push the genome past the error threshold (Fig. 3D). These simulations use a single parameter regime. A sensitivity analysis exploring the dependence of these results on the cheater advantage *s*, the number of within-cell replication rounds *τ*, the protocell population size *M*, and the copy number *n* is presented in the Supplementary Material (Fig. S1), and confirms that the qualitative result, pregenome collapse and post-genome stability, is robust across all parameter combinations tested.

## Discussion

The protocell is one of the most consequential innovations in the history of life. By enclosing a collection of catalytic molecules within a membrane, the protocell creates a higher-level unit on which natural selection can act: on the collective fitness of the ensemble, rather than the replicative success of any individual molecule. This is a key reason why cellular life has come to dominate the biosphere. But the cell alone is not sufficient. It provides a powerful mechanism enabling collective-level selection and adaptation, but it alone cannot durably drive an evolutionary transition to cell-level individuality. Cell-level individuality requires a singular genome and a ribosome to non-heritably amplify the information within it, which together imposes germ-soma division of labor at the molecular level: heritable information is confined to the genome, and the enzymatic workforce is rendered evolutionarily inert.

The pattern we have identified, the suppression of lower-level Darwinian dynamics through the segregation of heritable information into a germ line, is a hallmark of evolutionary transitions in individuality (24–26). In multicellular organisms, germ-soma specialization removes somatic cells from the heritable lineage, and cancer represents the pathological reassertion of somatic-level selection (14, 27). In eusocial colonies, sterile workers are de-Darwinized by the loss of direct reproduction (28). In the endosymbiotic origin of organelles, gene transfer to the host nucleus strips endosymbionts of their evolutionary autonomy (29). In each case, formerly independent Darwinian units are brought under the control of a higher-level individual (30). What we have argued is that the genome-plus-ribosome system is the earliest known instance of this logic. The central dogma (31) is a statement that information does not flow from protein back to nucleic acid: translation imposes a strong barrier to the reassertion of lower-level selection in the proteome, just as germ-line sequestration constrains somatic cell evolution in animals. Like other evolutionary transitions, this barrier is imperfect; prions partially re-Darwinize the soma, and phenomena such as RNA editing and translational recoding complicate the mapping between genome and proteome. But the core asymmetry, that no protein variant can rewrite its encoding gene, is shared across all cellular life, and this fact makes open-ended growth in cellular complexity possible. This cross-hierarchical pattern has been previously noted by Takeuchi and Kaneko (32), who generalize the central dogma as a division of labor between information transmission and expression recurring across biological scales.

It is worth contrasting our proposal with the hypercycle, the elegant framework introduced by Eigen and Schuster (16) to address the same information-capacity barrier. Eigen and Schuster recognized that the error threshold limits how much information any single self-replicative RNA can maintain, and that a translation system requires more information than one molecule can encode at early replication fidelities. Their solution was a cyclic catalytic linkage among multiple self-replicative RNA species, in which each member catalyzes the replication of the next. This arrangement allows cooperation among units that would otherwise be strict competitors, integrating their information content without placing it on a single molecule. The mathematical analysis shows that only cyclic linkages, not chains or branching networks, have the stability properties required for such integration. The hypercycle, unfortunately, does not solve the problem we have identified as the more fundamental constraint. The members of a hypercycle remain fully Darwinian: each replicates, accumulates mutations, and is subject to selection at the molecular level. In the language of Eq. 4, the hypercycle reorganizes the competitive relationships among replicative units but does not eliminate the within-protocell selection term. Mutation is still biased toward loss of function, less functional and parasitic variants can still arise and spread within the cycle, and these problems worsen as the number of cooperating species grows, for the same scaling reasons described above. The vulnerability of cooperative molecular networks to parasitic replicators was recognized early and has been observed experimentally in compartmentalized RNA replication systems (10–13). The genome-plus-ribosome architecture does something qualitatively different: it removes the molecular workforce from the heritable lineage entirely, setting the within-protocell selection term to zero rather than merely moderating it. It is the asymmetry of information flow in translation, not the topology of catalytic coupling, that breaks the evolutionary agency of enzymes and permits open-ended growth in network complexity.

A major open question that our framework does not resolve is how the translation system itself could have originated within the very environment we argue is hostile to increasing complexity. A primitive ribosome, however rudimentary, requires multiple coordinated components, and these components would have needed to evolve and be co-maintained in pre-genome protocells subject to exactly the coordination and conflict pressures we describe. This is a bootstrapping problem: the solution to the complexity barrier is itself complex. We do not attempt to resolve this here, but we note that the earliest proto-ribosome need not have resembled the modern translation apparatus. Structural analysis of ribosomal RNA across the tree of life suggests that the ribosome evolved by accretion, growing outward from a simple core (33). In this model, the peptidyl transferase center, the catalytic heart of the ribosome, was assembled from only a handful of small ancestral RNA expansion segments, and initially catalyzed only nonspecific, noncoded condensation of amino acids and possibly other substrates. Coded translation, decoding, and energy-driven translocation were not acquired until much later phases of ribosomal evolution. The earliest proto-ribosome, then, was not the elaborate molecular machine of modern biology but a small catalytic RNA, perhaps initially selected for some other function such as cofactor synthesis (34) or creation of short RNA-stablizing peptides (35), that provided enough of a separation between information and function to begin the process of de-Darwinization. Importantly, our simulations show that between-protocell selection can maintain collective function in networks of modest complexity (Fig. 3D), suggesting that the pre-genome architecture is not inherently doomed at the scale of a minimal proto-translation apparatus, even if it cannot sustain the open-ended growth in complexity that followed. Once even a partial germ-soma separation existed, it would have been self-reinforcing: protocells with better translation could tolerate greater network complexity, creating selection for improved translation, and additional structural elements could accrete onto the ribosome in step with this expanding capacity. Our model also assumes that the genome replicates once per protocell generation; the evolution of this coupling is itself nontrivial, because any linked molecule that replicates more slowly than its unlinked competitors faces a within-cell selective disadvantage. Experimental evolution of cooperative RNA replicators in compartments confirms this tension: physical linkage between genes evolved but did not dominate the population, because shorter unlinked fragments replicated faster (36).

The genome-plus-ribosome architecture is the fundamental innovation: it separates heritable information from the enzymatic workforce, solving both the coordination problem and the conflict problem as described above. The subsequent transition to DNA genomes extends this logic by molecularly imposing a second germ-soma specialization within the nucleic acid compartment itself. DNA replicates once per cell cycle and is read out unidirectionally into RNA, which is used to amplify genetic information for increased translation and degraded without self-replicating; RNA in cellular life is an informational dead end, just as proteins are. Two of the operational differences between DNA and RNA are chemical: the deoxyribose backbone confers greater resistance to hydrolysis (and reduced folding flexibility and catalytic potential), and the substitution of thymine for uracil enables enzymatic detection of cytosine deamination (37, 38). The third is entirely the result of evolution: DNA polymerases carry proof-reading exonuclease domains, whereas RNA polymerases generally do not. There is no structural necessity for this third asymmetry; evolution has simply invested in replication fidelity for the germ line and not for the soma. Together, these properties dramatically expand the maximum genome size that can be maintained below the error threshold: at per-nucleotide mutation rates typical of RNA replication (*µ*_*n*_ ≈ 10^−4^; (39)), viable genome size is limited to roughly 10^4^ nucleotides (15), whereas DNA, with mutation rates several orders of magnitude lower (40), raises this ceiling correspondingly. The germ-soma interpretation of the DNA/RNA boundary is reinforced by the pattern of its violation: just as prions partially re-Darwinize the proteome (20), reverse transcriptases allow RNA to write back into the DNA germ line and RNA-dependent RNA polymerases allow the replication of short autonomous RNA molecules, and the enzymes that catalyze these flows are overwhelmingly encoded by genomic parasites and viruses.

Protocells, an informational RNA genome, and the ribo-some are the trifecta which makes cellular life possible. The protocell provides compartmentalization and a unit of selection at the collective level; the genome consolidates heritable information into a single replicating molecule; and the ribosome translates that information into a functional, de-Darwinized workforce. Together, these three innovations make selection at the level of the cellular collective overwhelmingly effective, suppressing the within-cell evolutionary processes that would otherwise erode collective function. When combined with the higher fidelity of DNA, the amount of information stored in the genome can expand by orders of magnitude, underpinning the origin of life as we know it.

## Supporting information

Simulation scripts

## Acknowledgements

We thank Carl Simpson, Ryo Mizuuchi, Ryoka Kuwabara, Loren Williams and Nick Hud for helpful discussion and suggestions. This work was supported by The John Templeton Foundation Award 63580 to W.C.R.

