## Supplementary information for "De-Darwinizing the proteome: the genome as the original germ line"

**William C. Ratcliff**[1,✉]

[1]School of Biological Sciences, Georgia Institute of Technology, Atlanta, GA 30332, USA

### Sensitivity analysis of multilevel selection simulations

The main text presents simulations of pre-genome and post-genome protocells under a single parameter regime (mutation rate $\mu = 0.005$, cheater advantage $s = 0.02$, within-cell replication rounds $\tau = 10$, protocell population size $M = 100$, copies per ribozyme type $n = 10$, per-gene genome mutation rate $U = 0.005$, translational error rate $\mu_t = 0.005$). Here we explore the sensitivity of the central result, that pre-genome protocells lose collective function as network complexity $k$ grows while post-genome protocells remain stable, to systematic variation in each of these parameters. For each parameter scan, all other parameters are held at their default values. Equilibrium fitness is computed as the mean fitness across generations 200 to 300, averaged over 3 replicate simulations per condition (Fig. 1).

**Cheater replication advantage ($s$).** We varied the within-cell replication advantage of cheater variants from $s = 0$ (no cheaters, mutation only) to $s = 0.05$ (strong cheater advantage), at fixed $\mu = 0.005$, $\tau = 10$, $n = 10$, and $M = 100$ (Fig. 1A). Even in the absence of cheaters ($s = 0$), pre-genome protocells suffer progressive fitness decline as $k$ increases, because mutational degradation alone, operating through Muller's ratchet across $k$ independent ribozyme types, is sufficient to erode collective function. Introducing cheaters accelerates this decline substantially: at $s = 0.05$, pre-genome equilibrium fitness is effectively zero for $k$ greater than approximately 10. The post-genome architecture is entirely unaffected by the value of $s$, because proteins do not replicate and therefore cannot cheat. This result underscores the point made in the main text: even in the most favorable case for the pre-genome architecture (no cheaters at all), mutational degradation alone imposes a ceiling on network complexity. Cheaters make the problem worse, but they are not the root cause.

**Within-cell replication rounds ($\tau$).** We varied the number of ribozyme replication rounds per protocell generation from $\tau = 2$ to $\tau = 20$, at fixed $\mu = 0.005$, $s = 0.02$, $n = 10$, and $M = 100$ (Fig. 1B). The parameter $\tau$ controls how many rounds of within-cell Darwinian dynamics occur before the next protocell-level selection event. As predicted by the multilevel Price equation analysis in the main text, increasing $\tau$ amplifies the within-protocell selection term, because more rounds of replication allow more mutation accumulation and more cheater spread before between-protocell selection can act. At $\tau = 2$, pre-genome protocells retain modest fitness even at $k = 20$, because within-cell degradation is limited by the small number of replication rounds. At $\tau = 20$, the decline is severe and begins at very low $k$. The post-genome architecture is unaffected by $\tau$, since it has no within-cell replication dynamics to compound. This result highlights that the ratio of within-cell to between-cell evolutionary rates is a critical determinant of the pre-genome ceiling, and that reducing this ratio (for example, through faster protocell division relative to ribozyme replication) could in principle extend the range of viable complexity, though not eliminate the fundamental problem.

**Protocell population size ($M$).** We varied the number of competing protocells from $M = 20$ to $M = 200$, at fixed $\mu = 0.005$, $s = 0.02$, $\tau = 10$, and $n = 10$ (Fig. 1C). Larger populations provide more effective between-protocell selection, because the covariance between fitness and trait value ($\mathrm{Cov}(W_j, Z_j)$ in the Price equation) is estimated more precisely and extreme variants are more likely to be represented. We find that increasing $M$ from 20 to 200 provides a modest improvement in pre-genome equilibrium fitness, but the qualitative pattern is unchanged: pre-genome fitness still collapses as $k$ grows, regardless of population size. For the post-genome architecture, larger populations improve the efficiency of purifying selection against genomic mutations, resulting in slightly higher equilibrium fitness at large $k$. The insensitivity of the pre-genome collapse to population size is consistent with the argument that within-cell degradation, not the inefficiency of between-cell selection, is the binding constraint. Even a very large population of protocells cannot select effectively when within-cell dynamics have driven all protocells to similarly degraded states, leaving little between-protocell variance for selection to act upon.

**Ribozyme copy number ($n$).** We varied the number of copies per ribozyme type from $n = 5$ to $n = 40$, at fixed $\mu = 0.005$, $s = 0.02$, $\tau = 10$, and $M = 100$ (Fig. 1D). Counterintuitively, higher copy numbers accelerate the pre-genome fitness collapse rather than slowing it. This result can be understood through the lens of the multilevel Price equation, where the change in mean trait value depends on the

DRAFT

balance between a positive between-cell selection term and a negative within-cell selection term. Higher $n$ shifts both terms against the pre-genome architecture simultaneously.

First, increasing $n$ weakens between-protocell selection. The stochastic corrector model depends on binomial partitioning at cell division to regenerate compositional variance each generation, providing the raw material for protocell-level selection. The variance of the sampled fraction scales as $p(1-p)/n$, so increasing $n$ tightens the distribution of ribozyme compositions across the population. This reduces the covariance $\mathrm{Cov}(W_j, Z_j)$ in the between-cell selection term of the Price equation, leaving selection with less to act on.

Second, increasing $n$ strengthens within-cell selection for cheaters. In a small molecular population, genetic drift can counteract the deterministic replication advantage $s$ of cheater variants, occasionally pushing the functional fraction back up and slowing the rate at which cheaters spread. With larger $n$, the within-cell dynamics track the deterministic expectation more faithfully: cheaters reliably outcompete functional variants at rate $s$ per replication round, and mutation erodes function at rate $\mu$ per round, with little stochastic noise to disrupt either process. The within-cell selection term $E(W_j \cdot \Delta Z_j)$ therefore becomes more negative as $n$ increases.

The combined effect is striking. Higher copy number simultaneously weakens the positive term in the multilevel Price equation (less between-cell variance for selection to exploit) and strengthens the negative term (more efficient within-cell cheater spread), producing a faster collapse of collective function as $k$ grows. This result underscores the dependence of the pre-genome architecture on stochastic noise at multiple levels: noise at division to generate between-cell variance, and noise within cells to slow the deterministic spread of selfish variants. The post-genome architecture, by contrast, is independent of $n$, since copy number is a property of the ribozyme-based system and has no analog in the genome-plus-ribosome framework where proteins are produced fresh each generation from the genome.

**Summary.** These results confirm that the conclusions drawn in the main text are robust across a broad range of parameter values. The pre-genome collapse with increasing network complexity is a consistent feature of the architecture, not an artifact of a particular parameter choice. Varying the cheater advantage, the number of within-cell replication rounds, the protocell population size, and the ribozyme copy number all modulate the quantitative details of the collapse, shifting the critical $k$ at which fitness drops to negligible levels, but none of these variations rescues the pre-genome architecture at high complexity. The copy number result is particularly instructive: increasing $n$ might naively be expected to stabilize the pre-genome system by buffering against stochastic loss of functional variants, but instead it accelerates the collapse by undermining the very noise that the stochastic corrector mechanism depends on. The post-genome architecture, by contrast, is stable across the full range of conditions tested, consistent with the theoretical prediction that eliminating the within-protocell selection term from the Price equation produces a qualitatively different evolutionary regime.

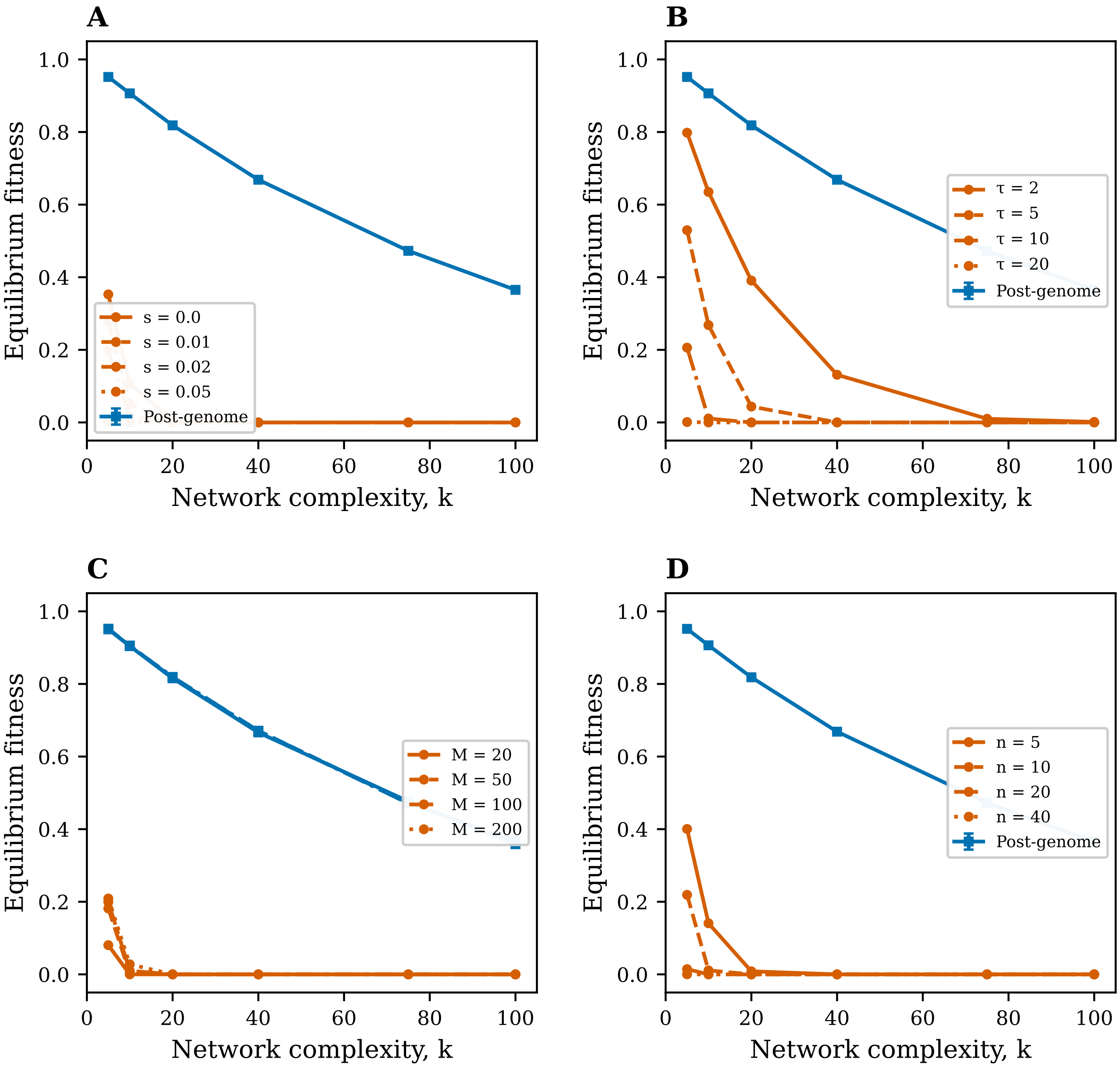


**Fig. 1. Sensitivity analysis of pre-genome and post-genome protocell fitness to key model parameters.** In all panels, orange curves show pre-genome equilibrium fitness and blue curves show post-genome equilibrium fitness as a function of network complexity $k$. Equilibrium fitness is averaged over generations 200–300 in populations of protocells with between-protocell selection. Default parameters unless otherwise varied: $\mu = 0.005$, $s = 0.02$, $\tau = 10$, $M = 100$, $n = 10$, $U = 0.005$, $\mu_t = 0.005$. (**A**) Varying cheater replication advantage $s$ from 0 (no cheaters) to 0.05 (strong cheater advantage). Pre-genome fitness declines with increasing $s$ at all values of $k$. Post-genome fitness is independent of $s$. (**B**) Varying within-cell replication rounds per protocell generation $\tau$ from 2 to 20. More within-cell replication rounds accelerate pre-genome fitness decline. Post-genome fitness is independent of $\tau$. (**C**) Varying protocell population size $M$ from 20 to 200. Larger populations provide modestly more effective between-protocell selection, but the qualitative pre-genome collapse is unchanged. Post-genome protocells benefit slightly from larger population size at high $k$. (**D**) Varying ribozyme copy number per type $n$ from 5 to 40. Higher copy numbers accelerate the pre-genome fitness collapse by simultaneously reducing between-cell variance (weakening protocell-level selection) and allowing more efficient within-cell cheater spread (strengthening within-cell selection). Post-genome fitness is independent of $n$. Error bars (where shown) represent standard deviation across 3 replicate simulations.
