## Supplementary material for "De-Darwinizing the proteome: the genome as the original germ line": Simulation scripts: make_figures-2.py

```
"""
Generate all figures for "The genome as germ line" paper.

Figure 1 (figure1.pdf): The coordination problem (binomial partitioning).
Figure 3 (figure2_merged.pdf): De-Darwinization and multilevel selection (4 panels).
  - A: Schematic fitness decay (pre-genome) vs. constant fitness (post-genome).
       Uses a rescaled effective decay rate for visualization; the unscaled
       parameters drive pre-genome fitness to zero within a few generations,
       which is shown explicitly in panels C and D.
  - B: Translational error cost as a function of network complexity.
  - C: Stochastic simulation with between-protocell selection, trajectories at k=10,20,50.
  - D: Equilibrium fitness vs. k for both architectures.

Note: Figure 1 in the paper (overview_figure.jpg) is created separately.

Usage:
    python make_figures.py
"""

import numpy as np
import matplotlib.pyplot as plt
import matplotlib as mpl

# ── Global style ──────────────────────────────────────────────────
mpl.rcParams.update({
    'font.family': 'serif',
    'font.size': 9,
    'axes.labelsize': 10,
    'axes.titlesize': 11,
    'legend.fontsize': 7,
    'xtick.labelsize': 8,
    'ytick.labelsize': 8,
    'lines.linewidth': 1.5,
    'figure.dpi': 300,
    'savefig.dpi': 300,
    'savefig.bbox': 'tight',
    'mathtext.default': 'regular',
})

colors = ['#2c3e50', '#c0392b', '#2980b9', '#27ae60', '#8e44ad']
pre_color = '#D55E00'   # orange (Okabe-Ito), colorblind-safe
post_color = '#0072B2'  # blue (Okabe-Ito), colorblind-safe


# ══════════════════════════════════════════════════════════════════
# SIMULATION FUNCTIONS
# ══════════════════════════════════════════════════════════════════

def simulate_pregenome(M, k, n, mu, s, tau, G, seed):
    """
    Simulate a population of M pre-genome protocells over G generations.

    Each protocell has k ribozyme types, n copies each.
    Within each protocell generation, tau rounds of ribozyme replication
    occur with:
      - mutation rate mu (functional -> non-functional, irreversible)
      - cheater replication advantage s

    Between-protocell selection: protocells reproduce proportional to
    fitness (product of functional fractions across all k types),
    maintaining constant population size M.

    Returns: 1D array of mean population fitness per generation.
    """
    rng = np.random.default_rng(seed)
    # x[j, i] = fraction of functional copies of type i in protocell j
    x = np.ones((M, k))
    mean_fitness = np.zeros(G)

    for gen in range(G):
        # Within-cell dynamics: tau rounds of replication
        for _ in range(tau):
            # Mutation: functional fraction decays by mu
            x = x * (1 - mu)
            # Cheater selection: non-functional variants replicate at rate 1+s
            denom = x + (1 - x) * (1 + s)
            x = x / denom

        # Stochastic sampling (finite copy number effects)
        x = rng.binomial(n, np.clip(x, 0, 1)).astype(float) / n

        # Protocell fitness: product over types
        fitness = np.prod(x, axis=1)
        mean_fitness[gen] = np.mean(fitness)

        # Between-protocell selection: resample proportional to fitness
        # Note: daughter protocells inherit parental ribozyme fractions
        # without binomial partitioning at division. This is a deliberately
        # favorable assumption for the pre-genome architecture (steelmanning),
        # isolating evolutionary erosion from the coordination problem.
        total_f = fitness.sum()
        if total_f > 0:
            probs = fitness / total_f
            parents = rng.choice(M, size=M, p=probs, replace=True)
            x = x[parents].copy()
        # else: population is extinct; x remains at zero

    return mean_fitness


def simulate_postgenome(M, k, U, mu_t, G, seed):
    """
    Simulate a population of M post-genome protocells over G generations.

    Each protocell has a genome of k genes.
    Per generation:
      - Each intact gene mutates (becomes non-functional) with prob U.
        Back-mutation is neglected, consistent with the analytical model.
      - Translational error rate mu_t: each intact gene contributes
        (1 - mu_t) to fitness; mutated genes contribute residual delta = 0.01.
      - Fitness = product of per-gene contributions.

    Between-protocell selection: protocells reproduce proportional to
    fitness, maintaining constant population size M. This is the sole
    mechanism removing deleterious mutations from the population (purifying
    selection at the protocell level).

    Returns: 1D array of mean population fitness per generation.
    """
    rng = np.random.default_rng(seed)
    # genome[j, i] = 1 if gene i in protocell j is intact, 0 if mutated
    genome = np.ones((M, k), dtype=np.int8)
    mean_fitness = np.zeros(G)
    delta = 0.01  # residual function of a mutated gene

    for gen in range(G):
        # Fitness: product of per-gene contributions
        gene_fit = genome * (1 - mu_t) + (1 - genome) * delta
        fitness = np.prod(gene_fit, axis=1)
        mean_fitness[gen] = np.mean(fitness)

        # Between-protocell selection (purifying selection at protocell level)
        total_f = fitness.sum()
        if total_f > 0:
            probs = fitness / total_f
            parents = rng.choice(M, size=M, p=probs, replace=True)
            genome = genome[parents].copy()

        # Forward genome mutation (irreversible)
        mutations = rng.random((M, k)) < U
        genome[mutations] = 0

    return mean_fitness


# ══════════════════════════════════════════════════════════════════
# FIGURE 1: THE COORDINATION PROBLEM
# ══════════════════════════════════════════════════════════════════

def make_figure1():
    fig, (ax_a, ax_b) = plt.subplots(1, 2, figsize=(6.5, 3.3))

    # Panel A: P_viable vs k for different n
    k_vals = np.arange(1, 102)
    for i, n in enumerate([3, 5, 8, 12, 20]):
        ax_a.plot(k_vals, (1 - 2**(1 - n))**k_vals,
                  color=colors[i], label=f'n = {n}')

    ax_a.set_xlabel('Network complexity, k')
    ax_a.set_ylabel(r'$P_{\mathrm{viable}}$')
    ax_a.set_xlim(1, 100)
    ax_a.set_ylim(0, 1.02)
    ax_a.legend(loc='center right', ncol=1,
                frameon=True, fancybox=True, framealpha=0.95,
                edgecolor='0.8', columnspacing=1.0, handlelength=1.5)
    ax_a.set_title('A', loc='left', fontweight='bold', pad=6)

    # Panel B: required n and total N to maintain P_viable >= 0.95
    k_range = np.arange(2, 201)
    n_req = 1 - np.log2(1 - 0.95**(1.0 / k_range))
    ax_b_tw = ax_b.twinx()
    ln1 = ax_b.plot(k_range, n_req, color=colors[0],
                    label='Copies per type, n')
    ln2 = ax_b_tw.plot(k_range, n_req * k_range, color=colors[1],
                       ls='--', label='Total molecules, N')
    ax_b.set_xlabel('Network complexity, k')
    ax_b.set_ylabel('Required copies per type, n', color=colors[0])
    ax_b_tw.set_ylabel('Total molecules, N = kn', color=colors[1])
    ax_b.tick_params(axis='y', labelcolor=colors[0])
    ax_b_tw.tick_params(axis='y', labelcolor=colors[1])
    ax_b.set_xlim(2, 200)
    ax_b.set_title('B', loc='left', fontweight='bold', pad=6)
    lns = ln1 + ln2
    ax_b.legend(lns, [l.get_label() for l in lns],
                loc='upper left', ncol=1,
                frameon=True, fancybox=True, framealpha=0.95,
                edgecolor='0.8', handlelength=1.5)

    fig.tight_layout()
    fig.savefig('figure1.pdf')
    plt.close(fig)
    print("Figure 1 (coordination problem) done.")


# ══════════════════════════════════════════════════════════════════
# FIGURE 3: DE-DARWINIZATION + MULTILEVEL SELECTION (merged A–D)
# ══════════════════════════════════════════════════════════════════

def make_figure3():
    # Simulation parameters
    M = 100       # protocell population size
    G = 500       # generations
    mu = 0.005    # mutation rate
    s = 0.02      # cheater replication advantage
    tau = 10      # within-cell replication rounds per protocell generation
    n = 10        # copies per ribozyme type
    U = 0.005     # per-gene genome mutation rate
    mu_t = 0.005  # translational error rate
    n_reps = 5    # replicate simulations

    fig, axes = plt.subplots(2, 2, figsize=(6.5, 6.2))
    ax_a, ax_b = axes[0]
    ax_c, ax_d = axes[1]

    # ── Panel A: schematic fitness decay (rescaled for visualization) ──
    # The actual decay rate k*mu*tau = 1.0 per protocell generation
    # would drive fitness to zero within a few generations and is not
    # informative as a continuous curve. We show a rescaled effective
    # rate (mu_eff = mu/100) so the qualitative shape of the decay is
    # visible across the 500-generation x-axis. The unscaled dynamics
    # are simulated stochastically in panels C and D.
    t = np.arange(0, 501)
    k_det = 20
    mu_eff = mu / 100  # illustrative effective rate, see caption
    s_eff = 0.02 / 100  # matching rescaling for the cheater curve
    ax_a.plot(t, np.exp(-k_det * mu_eff * tau * t),
              color=colors[1], label='Pre-genome (s = 0)')
    ax_a.plot(t, np.exp(-k_det * (mu_eff + s_eff) * tau * t),
              color=colors[4], ls='--',
              label='Pre-genome (s > 0, cheaters)')
    ax_a.plot(t, np.full_like(t, (1 - mu)**k_det, dtype=float),
              color=colors[3], lw=2, label='Post-genome')
    ax_a.set_xlabel('Protocell generations')
    ax_a.set_ylabel('Protocell fitness, W')
    ax_a.set_ylim(0, 1.05)
    ax_a.set_xlim(0, 500)
    ax_a.legend(loc='upper right', bbox_to_anchor=(1.0, 0.78),
                frameon=True, fancybox=True,
                framealpha=0.95, edgecolor='0.8', ncol=1)
    ax_a.set_title('A', loc='left', fontweight='bold', pad=6)

    # ── Panel B: translational error cost vs k ────────────────────
    k2 = np.arange(1, 502)
    for i, (mt, ml) in enumerate(zip(
            [1e-4, 1e-3, 1e-2, 5e-2],
            ['0.0001', '0.001', '0.01', '0.05'])):
        ax_b.plot(k2, (1 - mt)**k2, color=colors[i],
                  label=r'$\mu_t$ = ' + ml)
    ax_b.set_xlabel('Network complexity, k')
    ax_b.set_ylabel('Post-genome fitness')
    ax_b.set_xlim(1, 500)
    ax_b.set_ylim(0, 1.05)
    ax_b.legend(loc='upper right', frameon=True, fancybox=True,
                framealpha=0.95, edgecolor='0.8', ncol=1,
                columnspacing=1.0)
    ax_b.set_title('B', loc='left', fontweight='bold', pad=6)

    # ── Panel C: simulation trajectories at k = 10, 20, 50 ───────
    k_style = {10: '-', 20: '--', 50: ':'}
    for k_val in [10, 20, 50]:
        pre_runs = np.array([
            simulate_pregenome(M, k_val, n, mu, s, tau, G,
                               42 + r + k_val * 100)
            for r in range(n_reps)
        ])
        post_runs = np.array([
            simulate_postgenome(M, k_val, U, mu_t, G,
                                42 + r + k_val * 100 + 5000)
            for r in range(n_reps)
        ])
        ax_c.plot(np.arange(G), pre_runs.mean(0),
                  color=pre_color, ls=k_style[k_val], alpha=0.85,
                  label=f'Pre, k={k_val}')
        ax_c.plot(np.arange(G), post_runs.mean(0),
                  color=post_color, ls=k_style[k_val], alpha=0.85,
                  label=f'Post, k={k_val}')
        print(f'  Panel C: k={k_val} done')

    ax_c.set_xlabel('Protocell generations')
    ax_c.set_ylabel('Mean protocell fitness')
    ax_c.set_xlim(0, 500)
    ax_c.set_ylim(0, 1.05)
    ax_c.legend(loc='lower right', frameon=True, fancybox=True,
                framealpha=0.95, edgecolor='0.8', ncol=1,
                handlelength=1.8, fontsize=6.5)
    ax_c.set_title('C', loc='left', fontweight='bold', pad=6)

    # ── Panel D: equilibrium fitness vs k ─────────────────────────
    k_scan = np.array([5, 10, 15, 20, 30, 40, 50, 75, 100])
    pre_eq_m, pre_eq_s, post_eq_m, post_eq_s = [], [], [], []
    for k_val in k_scan:
        pre_eqs = [
            np.mean(simulate_pregenome(
                M, k_val, n, mu, s, tau, G,
                1000 + r + k_val * 100)[400:])
            for r in range(n_reps)
        ]
        post_eqs = [
            np.mean(simulate_postgenome(
                M, k_val, U, mu_t, G,
                1000 + r + k_val * 100 + 5000)[400:])
            for r in range(n_reps)
        ]
        pre_eq_m.append(np.mean(pre_eqs))
        pre_eq_s.append(np.std(pre_eqs))
        post_eq_m.append(np.mean(post_eqs))
        post_eq_s.append(np.std(post_eqs))
        print(f'  Panel D: k={k_val} done')

    ax_d.errorbar(k_scan, pre_eq_m, yerr=pre_eq_s, color=pre_color,
                  marker='o', markersize=4, capsize=3, label='Pre-genome')
    ax_d.errorbar(k_scan, post_eq_m, yerr=post_eq_s, color=post_color,
                  marker='s', markersize=4, capsize=3, label='Post-genome')
    ax_d.set_xlabel('Network complexity, k')
    ax_d.set_ylabel('Equilibrium fitness')
    ax_d.set_xlim(0, 105)
    ax_d.set_ylim(0, 1.05)
    ax_d.legend(loc='upper right', frameon=True, fancybox=True,
                framealpha=0.95, edgecolor='0.8', ncol=1,
                columnspacing=1.0)
    ax_d.set_title('D', loc='left', fontweight='bold', pad=6)

    fig.tight_layout(h_pad=2.0, w_pad=2.0)
    fig.savefig('figure2_merged.pdf')
    plt.close(fig)
    print("Figure 3 (de-Darwinization + multilevel selection) done.")


# ══════════════════════════════════════════════════════════════════
# MAIN
# ══════════════════════════════════════════════════════════════════

if __name__ == '__main__':
    make_figure1()
    make_figure3()
    print("\nAll figures complete.")
```
