## Supplementary material for "De-Darwinizing the proteome: the genome as the original germ line": Simulation scripts: make_supplement-2.py

```
"""
Supplementary Figure: Sensitivity analysis for pre-genome vs. post-genome protocells.

Explores how equilibrium fitness depends on:
  A: cheater advantage s (at fixed tau, mu)
  B: within-cell replication rounds tau (at fixed s, mu)
  C: protocell population size M
  D: ribozyme copy number n

Each panel shows equilibrium fitness (averaged over generations 400-500)
as a function of network complexity k, for several values of the
focal parameter, comparing pre-genome (orange) and post-genome (blue).

Usage:
    python make_supplement.py
"""

import numpy as np
import matplotlib.pyplot as plt
import matplotlib as mpl

# ── Global style (matches main figures) ──────────────────────────
mpl.rcParams.update({
    'font.family': 'serif',
    'font.size': 9,
    'axes.labelsize': 10,
    'axes.titlesize': 11,
    'legend.fontsize': 7,
    'xtick.labelsize': 8,
    'ytick.labelsize': 8,
    'lines.linewidth': 1.5,
    'figure.dpi': 300,
    'savefig.dpi': 300,
    'savefig.bbox': 'tight',
    'mathtext.default': 'regular',
})

pre_color = '#D55E00'   # orange (Okabe-Ito), colorblind-safe
post_color = '#0072B2'  # blue (Okabe-Ito), colorblind-safe


# ══════════════════════════════════════════════════════════════════
# SIMULATION FUNCTIONS (identical to main script)
# ══════════════════════════════════════════════════════════════════

def simulate_pregenome(M, k, n, mu, s, tau, G, seed):
    rng = np.random.default_rng(seed)
    x = np.ones((M, k))
    mean_fitness = np.zeros(G)
    for gen in range(G):
        for _ in range(tau):
            x = x * (1 - mu)
            denom = x + (1 - x) * (1 + s)
            x = x / denom
        x = rng.binomial(n, np.clip(x, 0, 1)).astype(float) / n
        fitness = np.prod(x, axis=1)
        mean_fitness[gen] = np.mean(fitness)
        total_f = fitness.sum()
        if total_f > 0:
            probs = fitness / total_f
            parents = rng.choice(M, size=M, p=probs, replace=True)
            x = x[parents].copy()
        # else: population is extinct; x remains at zero
    return mean_fitness


def simulate_postgenome(M, k, U, mu_t, G, seed):
    rng = np.random.default_rng(seed)
    genome = np.ones((M, k), dtype=np.int8)
    mean_fitness = np.zeros(G)
    delta = 0.01
    for gen in range(G):
        gene_fit = genome * (1 - mu_t) + (1 - genome) * delta
        fitness = np.prod(gene_fit, axis=1)
        mean_fitness[gen] = np.mean(fitness)
        total_f = fitness.sum()
        if total_f > 0:
            probs = fitness / total_f
            parents = rng.choice(M, size=M, p=probs, replace=True)
            genome = genome[parents].copy()
        mutations = rng.random((M, k)) < U
        genome[mutations] = 0
    return mean_fitness


def get_equilibrium(sim_func, args_list, n_reps=3, G=300, eq_start=200):
    """Run n_reps simulations and return mean and std of equilibrium fitness."""
    eqs = []
    for r in range(n_reps):
        args = args_list(r)
        trajectory = sim_func(*args)
        eqs.append(np.mean(trajectory[eq_start:]))
    return np.mean(eqs), np.std(eqs)


# ══════════════════════════════════════════════════════════════════
# SUPPLEMENT FIGURE
# ══════════════════════════════════════════════════════════════════

def make_supplement():
    fig, axes = plt.subplots(2, 2, figsize=(6.5, 6.2))
    ax_a, ax_b = axes[0]
    ax_c, ax_d = axes[1]

    k_scan = np.array([5, 10, 20, 40, 75, 100])
    G = 300
    n_reps = 3

    # Default parameters
    mu_default = 0.005
    s_default = 0.02
    tau_default = 10
    M_default = 100
    n_default = 10
    U_default = 0.005
    mu_t_default = 0.005

    # ── Panel A: varying cheater advantage s ──────────────────────
    s_vals = [0.0, 0.01, 0.02, 0.05]
    linestyles = ['-', '--', '-.', ':']

    for i, s_val in enumerate(s_vals):
        pre_m, pre_s_err = [], []
        post_m, post_s_err = [], []
        for k_val in k_scan:
            pm, ps = get_equilibrium(
                simulate_pregenome,
                lambda r, k=k_val, s=s_val: (
                    M_default, k, n_default, mu_default, s,
                    tau_default, G, 2000 + r + k * 100),
                n_reps, G)
            pre_m.append(pm)
            pre_s_err.append(ps)
            # Post-genome is independent of s
        # Post-genome only needs to be computed once (same for all s)
        if i == 0:
            for k_val in k_scan:
                pm, ps = get_equilibrium(
                    simulate_postgenome,
                    lambda r, k=k_val: (
                        M_default, k, U_default, mu_t_default, G,
                        3000 + r + k * 100),
                    n_reps, G)
                post_m.append(pm)
                post_s_err.append(ps)
            post_m_saved = post_m
            post_s_saved = post_s_err

        ax_a.plot(k_scan, pre_m, color=pre_color, ls=linestyles[i],
                  marker='o', markersize=3,
                  label=f's = {s_val}')
        print(f'  Panel A: s={s_val} done')

    ax_a.errorbar(k_scan, post_m_saved, yerr=post_s_saved,
                  color=post_color, marker='s', markersize=3,
                  capsize=2, label='Post-genome')
    ax_a.set_xlabel('Network complexity, k')
    ax_a.set_ylabel('Equilibrium fitness')
    ax_a.set_xlim(0, 105)
    ax_a.set_ylim(-0.05, 1.05)
    ax_a.legend(loc='lower left', frameon=True, facecolor='white', framealpha=0.95, ncol=1,
                handlelength=1.8, fontsize=6.5)
    ax_a.set_title('A', loc='left', fontweight='bold', pad=6)

    # ── Panel B: varying within-cell replication rounds tau ───────
    tau_vals = [2, 5, 10, 20]

    for i, tau_val in enumerate(tau_vals):
        pre_m, pre_s_err = [], []
        for k_val in k_scan:
            pm, ps = get_equilibrium(
                simulate_pregenome,
                lambda r, k=k_val, tau=tau_val: (
                    M_default, k, n_default, mu_default, s_default,
                    tau, G, 4000 + r + k * 100 + tau * 10000),
                n_reps, G)
            pre_m.append(pm)
            pre_s_err.append(ps)

        ax_b.plot(k_scan, pre_m, color=pre_color, ls=linestyles[i],
                  marker='o', markersize=3,
                  label=r'$\tau$ = ' + str(tau_val))
        print(f'  Panel B: tau={tau_val} done')

    ax_b.errorbar(k_scan, post_m_saved, yerr=post_s_saved,
                  color=post_color, marker='s', markersize=3,
                  capsize=2, label='Post-genome')
    ax_b.set_xlabel('Network complexity, k')
    ax_b.set_ylabel('Equilibrium fitness')
    ax_b.set_xlim(0, 105)
    ax_b.set_ylim(-0.05, 1.05)
    ax_b.legend(loc='center right', frameon=True, facecolor='white', framealpha=0.95, ncol=1,
                handlelength=1.8, fontsize=6.5)
    ax_b.set_title('B', loc='left', fontweight='bold', pad=6)

    # ── Panel C: varying protocell population size M ──────────────
    M_vals = [20, 50, 100, 200]

    for i, M_val in enumerate(M_vals):
        pre_m, pre_s_err = [], []
        post_m_c, post_s_c = [], []
        for k_val in k_scan:
            pm, ps = get_equilibrium(
                simulate_pregenome,
                lambda r, k=k_val, M=M_val: (
                    M, k, n_default, mu_default, s_default,
                    tau_default, G, 6000 + r + k * 100 + M * 1000),
                n_reps, G)
            pre_m.append(pm)
            pre_s_err.append(ps)

            pm, ps = get_equilibrium(
                simulate_postgenome,
                lambda r, k=k_val, M=M_val: (
                    M, k, U_default, mu_t_default, G,
                    7000 + r + k * 100 + M * 1000),
                n_reps, G)
            post_m_c.append(pm)
            post_s_c.append(ps)

        ax_c.plot(k_scan, pre_m, color=pre_color, ls=linestyles[i],
                  marker='o', markersize=3,
                  label=f'M = {M_val}')
        ax_c.plot(k_scan, post_m_c, color=post_color, ls=linestyles[i],
                  marker='s', markersize=3)
        print(f'  Panel C: M={M_val} done')

    ax_c.set_xlabel('Network complexity, k')
    ax_c.set_ylabel('Equilibrium fitness')
    ax_c.set_xlim(0, 105)
    ax_c.set_ylim(-0.05, 1.05)
    ax_c.legend(loc='center right', frameon=True, facecolor='white', framealpha=0.95, ncol=1,
                handlelength=1.8, fontsize=6.5)
    ax_c.set_title('C', loc='left', fontweight='bold', pad=6)

    # ── Panel D: varying copy number n ────────────────────────────
    n_vals = [5, 10, 20, 40]

    for i, n_val in enumerate(n_vals):
        pre_m, pre_s_err = [], []
        for k_val in k_scan:
            pm, ps = get_equilibrium(
                simulate_pregenome,
                lambda r, k=k_val, n=n_val: (
                    M_default, k, n, mu_default, s_default,
                    tau_default, G, 8000 + r + k * 100 + n * 10000),
                n_reps, G)
            pre_m.append(pm)
            pre_s_err.append(ps)

        ax_d.plot(k_scan, pre_m, color=pre_color, ls=linestyles[i],
                  marker='o', markersize=3,
                  label=f'n = {n_val}')
        print(f'  Panel D: n={n_val} done')

    ax_d.errorbar(k_scan, post_m_saved, yerr=post_s_saved,
                  color=post_color, marker='s', markersize=3,
                  capsize=2, label='Post-genome')
    ax_d.set_xlabel('Network complexity, k')
    ax_d.set_ylabel('Equilibrium fitness')
    ax_d.set_xlim(0, 105)
    ax_d.set_ylim(-0.05, 1.05)
    ax_d.legend(loc='center right', frameon=True, facecolor='white', framealpha=0.95, ncol=1,
                handlelength=1.8, fontsize=6.5)
    ax_d.set_title('D', loc='left', fontweight='bold', pad=6)

    fig.tight_layout(h_pad=2.0, w_pad=2.0)
    fig.savefig('supplement_sensitivity.pdf')
    plt.close(fig)
    print("Supplement figure (sensitivity analysis) done.")


if __name__ == '__main__':
    make_supplement()
```
